# Targeted inhibition of SCF^SKP2^ confers anti-tumor activities resulting in a survival benefit in osteosarcoma

**DOI:** 10.1101/2023.05.13.540637

**Authors:** Jichuan Wang, Alexander Ferrena, Swapnil Singh, Ranxin Zhang, Valentina Viscarret, Waleed Al-Hardan, Osama Aldahamsheh, Hasibagan Borjihan, Amit Singla, Simon Yaguare, Janet Tingling, Xiaolin Zi, Yungtai Lo, Richard Gorlick, Edward L. Schwartz, Hongling Zhao, Rui Yang, David S. Geller, Deyou Zheng, Bang H. Hoang

## Abstract

Osteosarcoma(OS) is a highly aggressive bone cancer for which treatment has remained essentially unchanged for decades. Although OS is characterized by extensive genomic heterogeneity and instability, *RB1* and *TP53* have been shown to be the most commonly inactivated tumor suppressors in OS. We previously generated a mouse model with a double knockout (DKO) of *Rb1* and *Trp53* within cells of the osteoblastic lineage, which largely recapitulates human OS with nearly complete penetrance. SKP2 is a repression target of pRb and serves as a substrate recruiting subunit of the SCF^SKP2^ complex. In addition, SKP2 plays a central role in regulating the cell cycle by ubiquitinating and promoting the degradation of p27. We previously reported the DKOAA transgenic model, which harbored a knock-in mutation in p27 that impaired its binding to SKP2. Here, we generated a novel *p53-Rb1-SKP2* triple-knockout model (TKO) to examine SKP2 function and its potential as a therapeutic target in OS. First, we observed that OS tumorigenesis was significantly delayed in TKO mice and their overall survival was markedly improved. In addition, the loss of *SKP2* also promoted an apoptotic microenvironment and reduced the stemness of DKO tumors. Furthermore, we found that small-molecule inhibitors of SKP2 exhibited anti-tumor activities in vivo and in OS organoids as well as synergistic effects when combined with a standard chemotherapeutic agent. Taken together, our results suggest that SKP2 inhibitors may reduce the stemness plasticity of OS and should be leveraged as next-generation adjuvants in this cancer.

## 1. Introduction

Osteosarcoma (OS) is the most common primary bone malignancy in children and adolescents, characterized by aggressive growth and frequent metastases. Despite an aggressive treatment protocol, including multi-agent cytotoxic chemotherapy and ablative surgery, the long-term survival of patients has remained unchanged for decades [1]. Patients who present with overt metastatic or relapsed diseases sustain a 5-year survival rate of less than 30%, illustrating the inadequacy of the existing treatment approach [2]. Unfortunately, efforts to modify or intensify the current chemotherapy have failed to improve outcomes [3]. New strategies, including immunotherapeutic approaches, have been disappointing to date [4]. At present, the treatment options for patients with either metastatic or relapsed OS are minimal.

OS is characterized by a high level of genomic heterogeneity and instability, coupled with the lack of a defining fusion gene product or mutation, which may account for the lack of a targeted therapeutic approach [5]. However, recent next-generation sequencing studies have revealed that OS tumors share specific common alterations that can be identified from their genomic landscape. While a single treatment for OS is unlikely, these findings offer new hope for a targeted genome-informed approach [6]. Given that *RB1* and *TP53* are the most common co-inactivated tumor suppressors in OS, it is likely that the loss of function of these genes is a critical failure of the anti-tumor mechanism in this cancer. Our previous study revealed that *RB1* and *TP53* was the only pair of genetic alterations co-occurring in OS with statistical significance across multiple datasets [7]. Since p53 and Rb1 cannot be reactivated or re-introduced into the patient’s tumor, their inactivation poses a formidable challenge, making OS very difficult to treat.

The tumor suppressor pRb regulates gene expression by binding and deactivating the E2F transcription factor, a major regulator of cell cycle-dependent gene expression[8]. The loss of pRb leads to elevated E2F transcriptional activities, compromising the ability of cells to exit the cell cycle and leading to a sustained proliferative state[9]. pRb also exerts a significant transcription-independent cell cycle control by regulating protein stability through the ubiquitin-proteasome pathway. Previous studies have shown that a deletion of *SKP2* effectively blocks tumorigenesis of pituitary and small cell lung cancer when *Rb1* and *Trp53* are co-inactivated [10]. As shown by our group and others, SKP2 is commonly over-expressed in *RB1*-inactivated malignancies like OS, and SKP2 knockdown suppresses the proliferation and invasion of established OS cell lines[10, 11]. Although these studies suggest that Skp2 promotes OS progression, results from established cell lines that have been passaged for many years may not accurately reflect the complex biology of OS. To better replicate the clinical scenario, we aim to investigate the SKP1-Cullin1-F-box (SCF)^SKP2^ complex using more robust preclinical models such as genetically engineered mouse models (GEMM) and organoids.

As an E3 ligase of the SCF complex, SKP2 exerts its oncogenic role by regulating cellular protein turnovers via the ubiquitin-proteasome degradation pathway[11-15]. Cyclin-dependent kinase (CDK) inhibitors such as p27 and p21 can be processed by SKP2 for ubiquitin-mediated degradation, which in turn promotes a cell cycle progression and tumorigenesis[14, 16, 17]. Disrupting the binding of SKP2 with p27 in OS leads to a stabilization of p27 and subsequent apoptosis, revealing that p27 plays a protective role in OS[18]. This safeguard mechanism of p27 has also been demonstrated in preclinical models of prostate and lung cancers in which *Rb1* and *Trp53* are co-inactivated[10, 19]. At present, the role of SKP2 in OS tumorigenesis and its therapeutic utility are not fully characterized. It is unknown whether inhibiting SKP2 can suppress OS tumorigenesis or lead to improved outcomes.

OS tumors contain a subpopulation of tumor-initiating cells (TICs) associated with chemoresistance and clinical relapses [20, 21]. Previously, we have shown that SKP2 plays a crucial role in maintaining a mesenchymal state and enhancing tumor-initiating properties in synovial sarcoma [22]. We recently showed that the interaction of SKP2 and its substrate p27 enhances OS cancer stemness[18]. Furthermore, SKP2 inhibitors have been found to restrict stem-like properties and potentiate the sensitivity to chemotherapy in other tumor models[23]. At present, the role of SKP2 in tumor-initiating properties in OS is still unexplored. Investigating whether SKP2 promotes OS stemness may lead to novel treatments to prevent relapses and alternative strategies for patients whose tumors are resistant to first-line chemotherapy.

In this study, on the basis of our Rb1/p53 double-knockout (DKO) mouse model (*Osx1-Cre;Rb1*^*lox/lox*^*;p53*^*lox/lox*^), we generated a novel GEMM with a triple knockout (TKO) of *Rb1, Tp53*, and *SKP2* (*Osx1-Cre;Rb1*^*lox/lox*^*;p53*^*lox/lox*^; *SKP2*^*-/-*^). Our previous DKOAA model (*Osx1-Cre;Rb1*^*lox/lox*^*;Trp5*3^lox/*lox*^*;p27*^*T187A/T187A*^) [24], which harbors a disrupted SKP2-p27 axis, was also used in this study to test the hypothesis that SKP2 promotes OS tumorigenesis comprehensively. The effects of an SKP2 deletion on the overall survival, cellular proliferation, apoptosis, and stemness were assessed *in vitro* and *in vivo*. Additionally, we established an organoid model of DKO OS as a potential high-throughput platform to test the efficacy of SKP2 inhibitors. Finally, the potential synergy between SKP2 inhibitors and conventional chemotherapy for OS was examined using multiple algorithms.

## 2. Materials and methods

### Expression analysis of human OS from public online databases

RNA sequencing data of OS patients were obtained from the Pediatric Preclinical Testing Consortium (PPTC) [25, 26]. The normalized counts of *TP53, RB1* and *SKP2* gene expression from each patient were compared. Corresponding tumor samples were further subjected to protein extraction and detection of p53, Rb, and SKP2 protein levels. GAPDH was used as the internal control.

RNA sequencing and clinical data were accessed from the NCI TARGET Osteosarcoma study[27]. The Reactome Apoptosis gene signature of 179 apoptosis-related genes was accessed from the Molecular Signature Database via the R package msigdbr (v7.4.1), followed by a composite expression module score calculation[28]. A survival analysis was performed in R (v4.1.0) using the survival (v3.2.13) and survminer (v0.4.9) packages. To examine associations of composite expression module scores with survival in OS patients. The module score was analysed as a dichotomized variable at the median. The score was also analysed as a continuous variable in univariable and multivariable Cox proportional hazards models adjusted for metastatic status at diagnosis. The proportional hazards assumption was checked via an inspection of Schoenfeld residuals.

### Establishment of animal models

*Osx1-Cre* mice, *Rb1*^*lox/lox*^ mice, *Trp53*^*lox/lox*^ mice, *p27*^*T187A/T187A*^ mice and *SKP2*^*-/-*^ mice were described previously[10, 18]. First, *Rb1*^*lox/lox*^ mice were crossed with *Trp53*^*lox/lox*^ mice to generate *Trp53*^*lox/lox*^, *Rb1*^*lox/lox*^, mice, and were further crossed with *Osx1-Cre* mice to generate *Osx1-Cre; Trp53*^*lox/lox*^, *Rb1*^*lox/lox*^ mice. The *SKP2*^*-/-*^ mice were crossed with *Osx1-Cre*;*Trp53*^*lox/lox*^,*Rb1*^*lox/lox*^ mice to generate *Osx1-Cre;Rb1*^*lox/lox*^*;Trp5*3^lox/*lox*^; *SKP2*^*-/-*^ mice. The genotyping results of each step were shown (Figure S1), and the correct genotypes were selected in a stepwise manner. Exons 2-10 in Trp53, exon 3 in Rb1, and exon 1 in SKP2 were deleted (Figure S2). Animals were maintained under pathogen-free conditions in the Albert Einstein College of Medicine animal facility. Animal experimental protocols were reviewed and approved by the Animal Care and Use Committee (#20180401), conforming to accepted standards of humane animal care. The tumor diameter was measured every three days using a caliper, and the relative tumor volume was calculated by the following formula: (length x width^2^) x 0.526. The growth of limb tumors was assessed when tumor volumes reached approximately 500 mm^3^.

### Immunohistochemistry, immunofluorescence, and TUNEL assay

Tissue sections and staining methods have been described previously [16, 18]. For immunohistochemical (IHC) staining, the following antibodies were used: PCNA (Vector Labs, SP6), p27 (BD Biosciences, #610242), Cleaved caspase 3 (Cell Signaling Technology, #9661), Anti-alkaline phosphatase (Abcam, ab354), Anti-osteocalcin (Abcam, ab13420).TUNEL staining was performed with an Apoptosis Detection Kit (Millipore, S7100). Images were visualized with the EVOS® FL Auto Imaging System (AMC1000, ThermoFisher). For the staining quantification, images were taken under a 60X magnification. Approximately 300 cells in each image and 3 images from each animal were counted. Data were presented as means of 3 animals in each group.

### Establishment of mOS organoid and viability assay

Primary tumors from DKO and TKO animals were dissociated into single-cell suspensions and 3000 cells were embedded in growth factor-reduced Matrigel (Corning,#356255). Organoid expansion and viability assays were performed in DMEM/F12 medium containing mitogens (EGF, FGF2, and FGF10), R-Spondin (Wnt/b-catenin signaling agonist), Noggin and A83-01 (TGFβ inhibitors), and Y-27632 (Rock inhibitor). Organoids were cultured for up to 4 weeks for viability measurement by tracking individual organoids using EVOS® FL Auto Imaging System (AMC1000, ThermoFisher). Proliferation assays were performed as previously described [18, 22]. For drug testing, cells were dissociated from mouse tumors and cultured for six days for organoid formation. Organoids were then treated with drugs for the indicated duration and concentration. A quantification of organoid viability was performed using CellTiter-Glo 3D (Promega, #G9682) according to the manufacturer’s protocol. All experiments were done in triplets for each organoid, compound, and concentration. For immunohistochemical staining, organoids were harvested and embedded in HistoGel™ (Thermo, #HG4000012), followed by fixation in a 4% paraformaldehyde (PFA) solution in PBS overnight. Thereafter, organoids were embedded in paraffin and immunostainings were performed as previously described [16, 18].

### Cell culture, cell proliferation, and extreme limiting dilution analysis (ELDA)

Primary cells were prepared from mouse tumor tissues, which were minced and dissociated with collagenase II in EMEM for 30-60 minutes at 37°C

Proliferation assays were then performed as previously described [18, 22]. C1 and Pevonedistat were prepared as previously described[18]. Cells were plated overnight and treated with each compound for the indicated duration and concentration. ELDA was performed and analyzed based on the protocol from Hu et al. [29]. Cells were seeded in ultra-low attachment 96-well plates at decreasing densities (at 20, 10, 5, 1 cell/well, 24 duplicates per density) and cultured for 12 days. The spheroid formation was monitored starting from day three with the criteria of one or more spheres greater than ∼50 μm diameter. Representative spheres are shown in Figure 4C.

### Western blot analysis and qRT-PCR

Western blot and qRT-PCR experiments were performed as previously described [11]. Antibodies for Western blots included p27 (BD Bioscience, #610242), SKP2 (Proteintech, #15010-1-AP), p21 (Santa Cruz, #SC-397), and cleaved caspase 3 (#9661),α-tubulin (#2125), p53(#2524), SKP2 (#2652), Rb (#9309), GAPDH (#5174) were from Cell Signaling Technology. PCR primers were summarized in Supplementary Table 2.

### Flow cytometry for apoptosis, ALDH assay, cell cycle assay, CD117 and Prominin-1 staining

An Annexin V-APC/7-AAD Apoptosis Kit (Abnova, #KA3808) was used for the apoptosis analysis as described previously [18]. ALDH assays were performed as described previously [22]. Cell cycle assays were performed as described, and propidium iodide(30 μg/ml) was used as DNA content dye [16, 30]. For CD117 staining, 1×10^6^ primary culture tumor cells were stained based on a protocol recommended by the manufacturer. The anti-mouse CD117-PE(Miltenyi Biotec, #REA791) antibody was used. The REA control-PE, (Miltenyi Biotec, # REA293) was used as a background control. A BD Accuri C6 flow cytometer (BD Biosciences, San Jose, CA) was used for the analysis. Results were analyzed by the Flow Jo software (Version 10.1, OR, USA). All experiments were performed in triplicates.

### Drug synergy analysis

The quantification of cell viability after drug treatments was determined by the MTT method as previously described[18]. Synergistic effects of doxorubicin in combination with C1 or Pevonendistat in primary cultured cells were determined using SynergyFinder 2.0 [31]. This tool applies three well-known synergistic analysis reference models: The zero interaction potency (ZIP) model[32], the highest single agent (HSA) model [33], and the Bliss independence model[34]. Synergy landscape plots over the dose matrix were generated by SynergyFinder. In general, positive, zero, and negative synergy scores indicate synergy, additivity, and antagonism, respectively.

### Inducible SKP2 knockdown in primary mOS cells

A doxycycline-inducible knockdown of SKP2 was performed with a pTripZ lentiviral shRNA obtained from Dharmacon as described previously[10].

Specifically, the shRNA sequence for SKP2 knockdown in mouse OS cells was 5′-183 GCAAGACTTCTGAACTGCTAT-3′. Lentiviral helper constructs psPAX2 and pMD2.G (Addgene plasmid #12260 and #12259) were used. An empty vector was used as a control. For a lentivirus infection, cells were placed in 1 ml of virus-containing media with 8 µg/ml polybrene and spun at 1000 x g for 30-60 minutes at 32°C. A puromycin selection was then performed, followed by mRNA (RT-qPCR) and protein (western blot) measurements. Doxycycline (2 µg/ml) was used to induce SKP2 knockdown in cultured cells.

### Establishment and treatment of mOS xenograft models

2.5 × 10^6^ primary tumor cells were mixed in a 1:1 culture medium and Matrigel mixture (LEDV-free; Corning) mixture and injected subcutaneously into NSG mice (6–8 weeks). The tumor diameter was measured using calipers and the tumor volume was calculated as (length x width^2^) x 0.526. Drug treatments were started once tumors reached approximately 150-250 mm^3^. Animals were randomly assigned into control and treatment groups with C1 (50 mg/kg/day sc) or Pevonedistat (90 mg/kg/day sc). Drugs were dissolved in 0.5% w/v CMC containing 10% DMSO, 30% PEG300, and 5% Tween-80. The vehicle solvent was administered in the control group.

### Statistical Analyses

Survival curves were generated by the Kaplan-Meier method and compared between groups using the log-rank test. Differences between groups were assessed using the Student’s t-test for continuous variables and the Fisher’s exact test for categorical variables. Differences in proliferation, viability, and drug inhibition assays were analyzed by two-way ANOVA. All statistical analyses were performed using SPSS version 22 (IBM, Armonk, NY). All tests were two-sided, and a P-value < 0.05 was considered statistically significant.

## 3. Results

### 3.1 The loss of *RB1* and *TP53* and the gain of *SKP2* are common in OS

We previously reported that SKP2 is overexpressed in human OS[11]. In addition, the co-occurrence of *RB1* and *TP53* inactivation was statistically significant in human OS [18]. To further determine the relation between *RB1, TP53*, and *SKP2* in human OS, transcriptional expression of *RB1, TP53*, and *SKP2* were queried from Pediatric Preclinical Testing Consortium databases [25, 26]. RNA-seq data of 25 OS patients’ tumor samples were obtained. Normalized counts of *RB1, TP53*, and *SKP2* from transcriptome sequencing were used to determine the relative gene expression. The relative expression of *RB1, TP53* and *SKP2* was analyzed and plotted separately (Figure 1A-C). A few OS samples that expressed relatively high levels of *RB1* and *TP53* mRNA showed lower levels of *SKP2* mRNA (OS-49-STTX-sc, OS-46-AOX). In contrast, the majority OS samples (OS-42, OS-47-SJ, OS-52-SJ, OS-54-SJ, OS-2, OS-34-SJ, etc.) retained low or undetectable *RB1* and *TP53* mRNA levels but high levels of *SKP2* mRNA (Figure 1A-C). The protein expression pattern similarly showed a negative relationship between Rb/p53 and SKP2 (Fig. 1D-E). Compared to non-tumor mesenchymal cell lines (MSC, NDHF, NHOst), most OS samples showed strong SKP2 expression accompanied by low or undetectable Rb and p53 levels.

**Figure 1.**
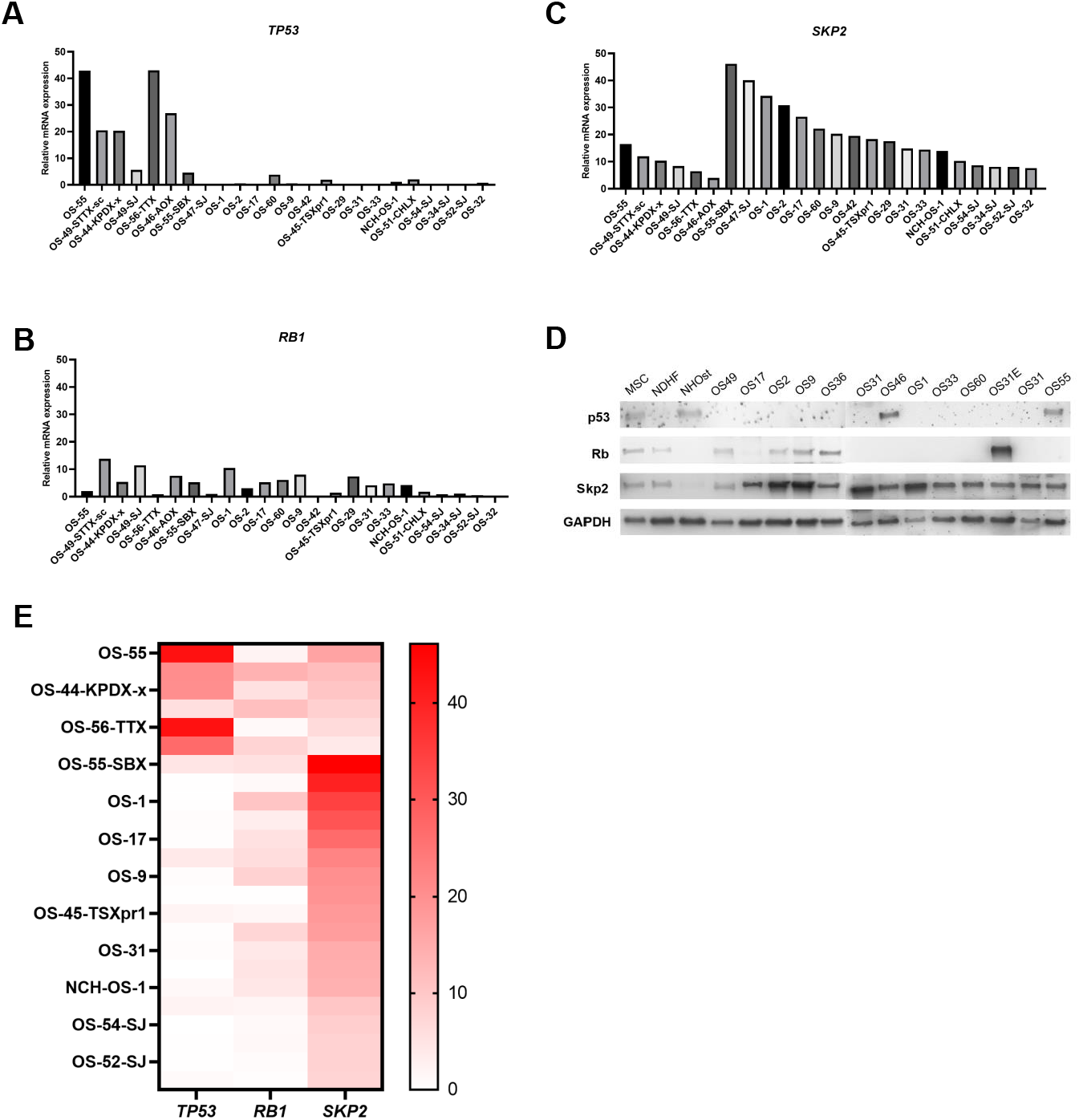
A loss of Rb and p53 and high SKP2 expression in patient-derived OS samples. A, Relative mRNA expression levels of *TP53, RB1* (B) and *SKP2* (C) as measured by RNA-seq in 24 PPTC osteosarcoma tumor samples. Relative mRNA expression levels were defined as counts. D. Western blot analysis for SKP2, TP53 and RB1 in protein lysates of the same OS samples as above. Three non-tumor mesenchymal cell lines, MSC (mesenchymal stem cells), NDHF (non-dermal human fibroblasts), NHOst (normal human osteoblasts), and 13 OS samples were included. GAPDH was used as a loading control. E. Heatmap of relative mRNA expression levels of *TP53, RB1* and *SKP2* as measured by RNA-seq in PPTC osteosarcoma tumor samples.

### 3.2 The loss of *SKP2* significantly delays the progression of OS

The OS tumorigenesis in DKO animals is invasive, metastatic, and lethal within 20–35 weeks. DKOAA and TKO mice are fertile without evidence of delayed or stunted growth. OS tumors developed with complete penetrance at an average of 154.65±29.08 days in DKO and 169±43.69 days in DKOAA mice. In contrast, TKO mice developed tumors at a much older age (251.86±48.53 days) (p<0.0001) (Figure 2G). Using a Kaplan-Meier analysis, the TKO mice (45.78±2.39 weeks, p<0.001) had significantly longer mean survival time than the DKO mice (32.57±1.32 weeks, p<0.001) and DKOAA (37.14±1.35 weeks, p<0.001) mice (Figure 2A). When DKO and TKO tumors were compared by IHC staining, an increase in p27 was observed in TKO tumors as a result of SKP2 deletion (Figure 2B). PCNA staining was reduced in TKO tumors when compared to DKO tumors (Figure 2B), indicating that SKP2 deletion suppresses tumor proliferation.

**Figure 2.**
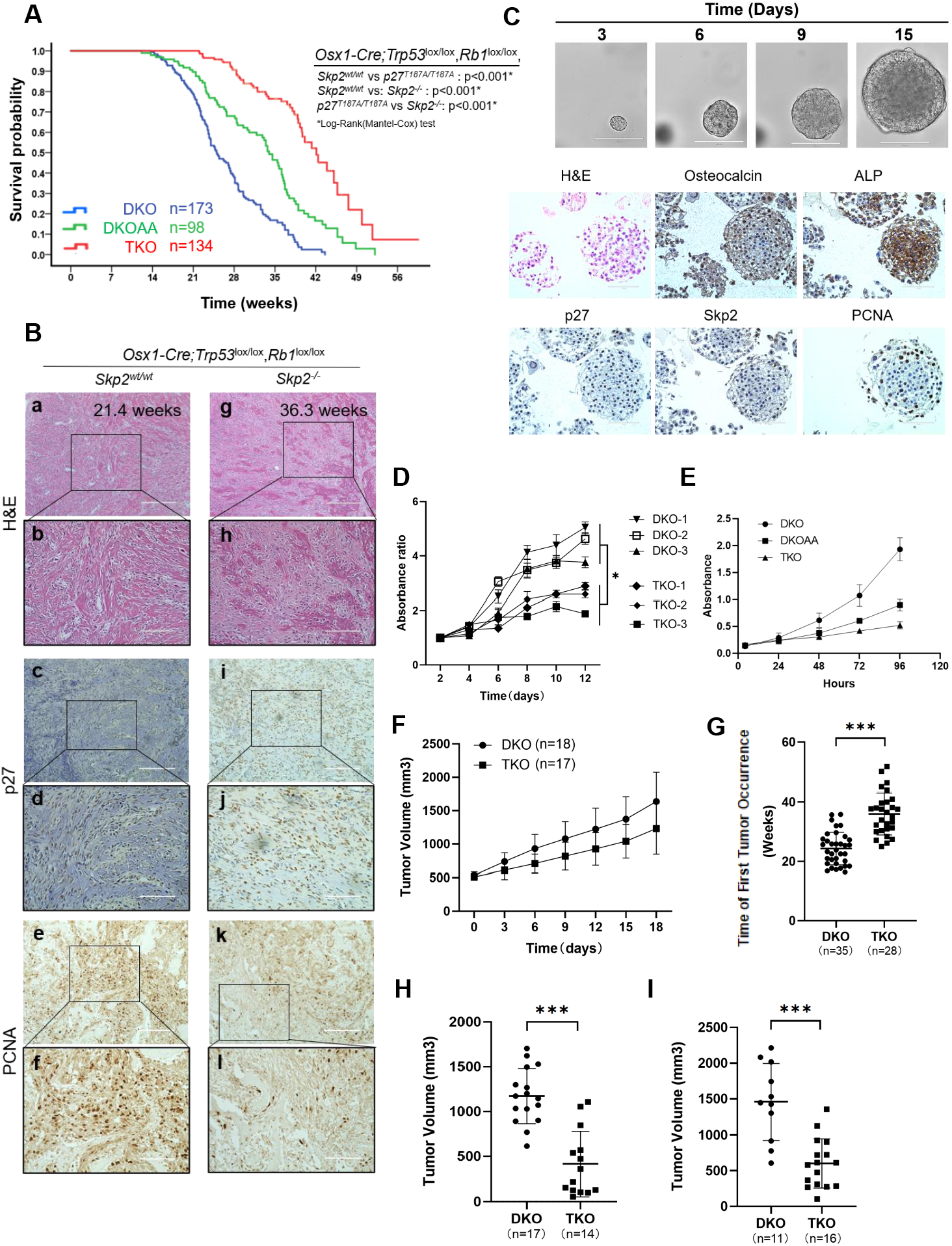
SKP2 deletion slows the progression of DKO OS tumorigenesis. (A) Kaplan-Meier survival analysis comparing the DKO, DKOAA and TKO cohorts of mice undergoing tumorigenesis. P-value is by a log-rank test and indicated in the figure. (B) Representative H&E, immunostaining of p27 and PCNA, of mOS tissue from the indicated genotype and age. Scale bars = 200 μm (a,c,e,g,i,k) and 100 μm (b,d,f,h,j,l). (C) DKO mOS tumor tissue-derived organoid culture. Upper figures indicate the proliferation of one representative organoid in the process of 15 days (Scale bars = 200 μm). Lower figures showed representative H&E of mOS organoid, and osteocalcin, alkaline phosphatase (ALP), p27, SKP2, and PCNA by immunohistochemistry. (D) Three independent mOS derived tumor organoids from DKO (DKO-1, DKO-2, and DKO-3) and TKO (TKO-1, TKO-2, and TKO-3) were cultured for the indicated duration, and relative proliferation rates were determined by the CellTiter-Glo method (*P*=0.035). (E) Graphs showing the proliferation of DKO, DKOAA and TKO tumor cells in monolayer cultures (*P*=0.001 DKO vs. DKOAA, *P*<0.001 DKO vs. TKO, *P*=0.006 DKOAA vs. TKO). (F) Mice with limb tumors were randomly selected from each group (n=18 from DKO group and n=17 from TKO group) once the tumor reached a volume around 500mm^3^, and measured every three days. The tumor growth rate was plotted and compared among the two groups. (G) The age of mice was documented and compared among the two groups when palpable or visible limb tumors were first discovered (*P*<0.001). Mice with measurable limb tumors at the same age (H) 28 weeks old, (I) 32 weeks old were randomly selected from each group, and tumor volume was measured and compared among two groups (*P*<0.001 and *P*<0.001). Error bars are SEM. ∗P< 0.05, ∗∗P< 0.01, and ∗∗∗P< 0.001.

To assess the tumor growth rate, animals with limb tumors were randomly selected from each group once the tumor volume reached approximately 500mm3. Tumor volumes were measured every three days and followed to the endpoint. The in vivo tumor proliferation was visibly slower in TKO mice than in DKO (p=0.19) (Figure 2F). When tumor volumes were compared at the same age, tumors from the DKO group were significantly larger than those from the TKO group at 28.5 weeks (1096.86 ± 267.04 mm^3^ vs. 804.11 ± 203.36 mm^3^, p=0.003) (Figure 2H) and at 32 weeks (1915.01 ± 728.62 mm^3^ vs. 1316.14 ± 623.16 mm^3^, p=0.039) (Figure 2I), respectively. *In vitro*, TKO tumor cells proliferated significantly less than both DKOAA (p<0.001) and DKO cells (p<0.001) (Figure 2E).

As a surrogate for in vivo tumor growth, both DKO and TKO organoid cultures successfully sustained and propagated from primary mOS tissues. One key features of organoids is it can be expanded and cryopreserved for organoid biobanking, thawed and rebuilt from cryopreservation as well easily. The reconstructed organoid recaptulate the morphology and progression of primary mOS tissues. A representative reconstructed DKO organoid was shown to proliferate for more than 2 months, with the diameter from less than 50 μm at day 3 to more than 100 μm at day 15 (Figure 2C). To compare the morphological and histological features of DKO organoids with their parental primary mOS tissues, we performed H&E staining and immunohistochemistry (IHC) analysis. Similar to the character of original mOS tissues, the H&E staining images of organoids exhibited cellular morphology with the obvious nucleoli and cytoplasmic content [18]. The IHC staining revealed that the organoids retained the high osteocalcin and alkaline phosphatase expression, both were consided as OS markers and similarly expressed on their parental mOS tissues. The similar pattern of relative high Skp2 and low p27 also exhibit the tumor feature compared with DKO mOS primary tissue. The positive staining of PCNA revealed the maintenance of the proliferation (Figure 2C). These data indicated that mOS organoids were stable to recapitulate the histology of parental DKO tumors. When tumors from three individual DKO and TKO mice were monitored for 2 weeks, the growth rate of the TKO group was significantly slower than that of the DKO cohort (p<0.001) (Figure 2D). Together, these data suggest that the SKP2 deletion significantly delays the occurrence and progression of DKO tumors and extends the survival of DKO mice.

### 3.3 A deletion or knockdown of SKP2 in OS stabilizes p27 and E2f1 apoptotic-related targets and leads to increased apoptosis

To determine whether an SKP2 deletion leads to an accumulation of p27, the histological features of DKO and TKO tumors were compared. By immunohistochemical (IHC) staining, p27 protein levels were markedly elevated in TKO compared to those in DKO tumors (Figure 2B). Western blots of tumor lysates from four animals in each group showed elevated p27 protein levels with corresponding decreases in SKP2 (Figure 3G). TUNEL and cleaved caspase-3 immunostainings indicated that TKO tumors sustained more apoptosis than DKO tumors (Figure 3A). Quantification of TUNEL staining also confirmed that the TKO cohort exhibited increased apoptosis <u>(p<0.001)</u> (Figure 3B). A flow cytometric analysis using Annexin V/7-AAD showed a significant increase in the early apoptotic population in TKO cells <u>(p<0.001)</u> (Figure 3E). PI staining showed a significant increase in cellular fractions later than G2 (1.9% vs. 14.2%), suggesting that TKO cells continued accumulating DNA content without dividing, decreasing proliferation (Figure 3F).

**Figure 3.**
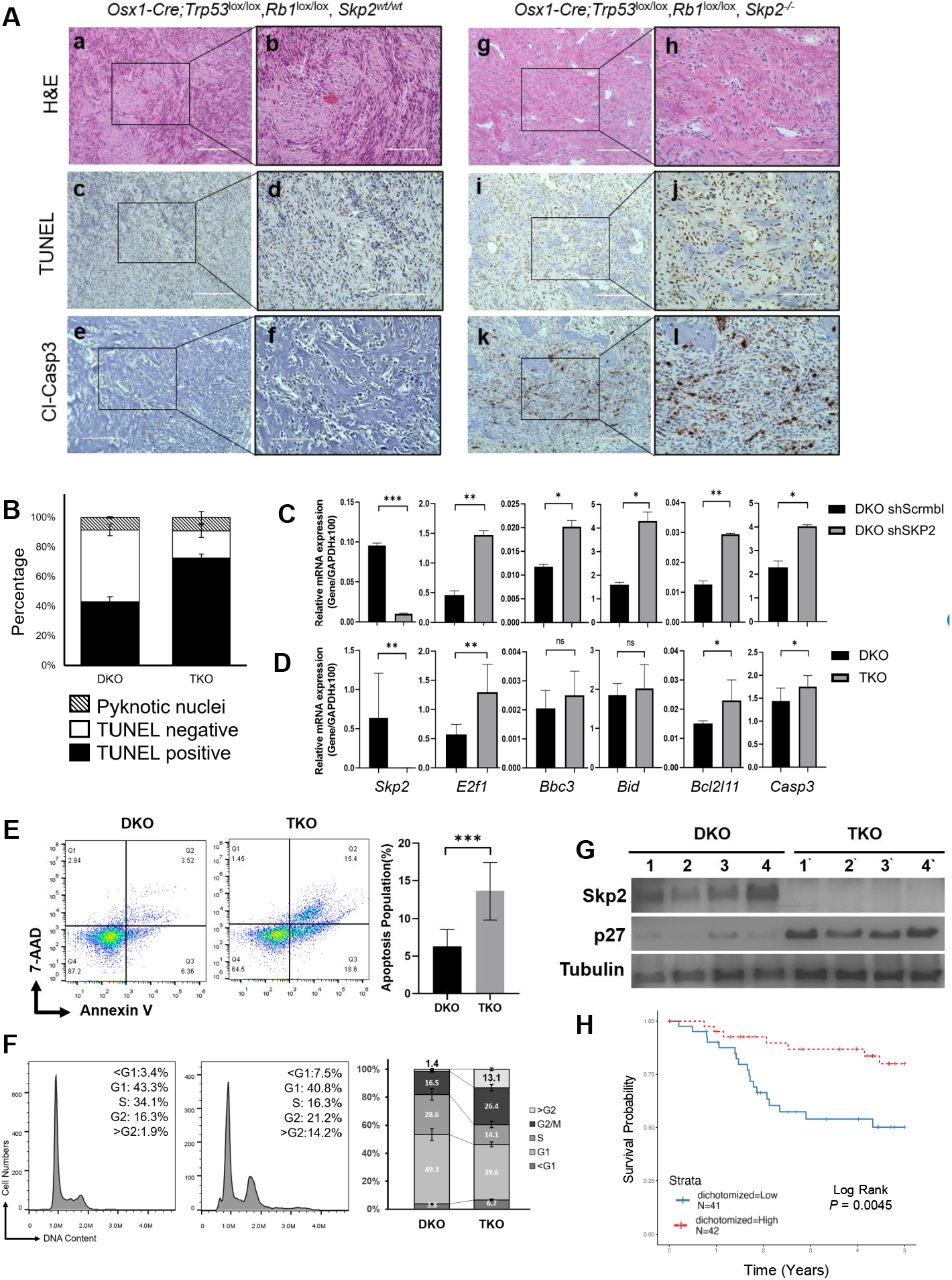
*SKP2* deletion increases apoptosis in DKO OS tumors. (A) Consecutive OS tumor sections of each genotype, as indicated, stained with H&E, TUNEL, and immunostaining of cleaved caspase-3. Scale bars = 200 μm (a, c, e, g, i, and k) and 100 μm (b, d, f, h, j, and l). (B) Quantification of apoptosis by TUNEL staining. Arrows demonstrate how total apoptosis was quantified. The bar graph was based on the average quantification of apoptosis from three mice. (C) SKP2 knockdown was achieved by shRNA and validated by qRT-PCR in DKO cells, followed by detection of mRNA levels of genes (*SKP2, E2f1, Bbc3, Bid, Bcl2l11, Casp3*) relative to GAPDH. (D) DKO and TKO cells were subject to qRT-PCR to determine mRNA levels of the same genes relative to GAPDH. (E) 7-exceAAD and annexin V staining of tumor cells of the indicated genotypes. Flow cytometry was used to detect and quantify apoptosis in the sub-G1 population. Error bars indicate SEM of the means of three samples. (F) Propidium iodide-based DNA content FACS (representative of three experiments) to detect and quantify cell phases in the population, as marked and plotted on the right. (G) Protein levels were determined by western blotting of OS tissues from the indicated genotypes. (H) Kaplan-Meier plot comparing survival of NCI TARGET OS cohort stratified by dichotomized expression of the Reactome Apoptosis gene signature at the median. Red line indicates patients overexpressing genes associated with apoptosis. Error bars are SEM. ∗P< 0.05, ∗∗P< 0.01, and ∗∗∗P< 0.001.

To elucidate the mechanistic consequences of an SKP2 inhibition, we examined the expression of *E2f1* and its apoptotic-related targets. SKP2-knockdown DKO cells (DKO shSKP2) (Figure 3C) and TKO cells (Figure 3D) were analyzed. *E2f1* mRNA levels were consistently elevated in DKO shSKP2 (p=0.005) and TKO (p=0.0013) groups compared to t he DKO scrambled control. Apoptotic-related targets of *E2f1* (*Bbc3, Bid, Bcl2l11*, and *Casp3*) were also elevated. Together, these data suggest that SKP2 deletion promotes a pro-apoptotic microenvironment in OS tumors in the absence of p53 and Rb1.

To further examine the contribution of apoptotic genes in human OS, we examined a cohort of 89 patients from the NCI TARGET database[27]. The overexpression of a composite expression score of the 179 apoptotic genes was found to be significantly correlated with improved survival in OS patients (p=0.0045) (Figure 3H, Table S3).

### 3.4 SKP2 inhibition reduces tumor-initiating properties of DKO tumors

Previously we reported that the binding of SKP2 to p27 promotes the stemness properties of OS [18]. To further elucidate the effects of inhibiting SKP2, mRNA levels of several OS stemness markers (*Aldh1a1, Aldh2, Aldh7a1, Kit*, and *Prom1*) were probed in DKO cells after SKP2 knockdown (Figure 4A) and in TKO cells (Figure 4B). We observed that stemness markers were down-regulated consistently in both groups. To further assess cancer stemness, a FACS analysis was performed, which demonstrated that TKO cells expressed significantly less CD117 than DKO cells (Figure 4E) (p<0.001). To determine whether this marker expression correlated with a stemness phenotype, ALDH activity assays were performed, which revealed a lower proportion of ALDH^high^ cells in the TKO group (p<0.001) (Figure 4D). Furthermore, an ELDA analysis revealed a lower stem cell frequency and self-renewal capacity (p<0.001) in TKO cells compared to those in DKO (Figure 4C). Overall, these results strongly suggest that SKP2 plays a crucial role in promoting tumor-initiating properties of OS when p53 and pRb are co-inactivated.

**Figure 4.**
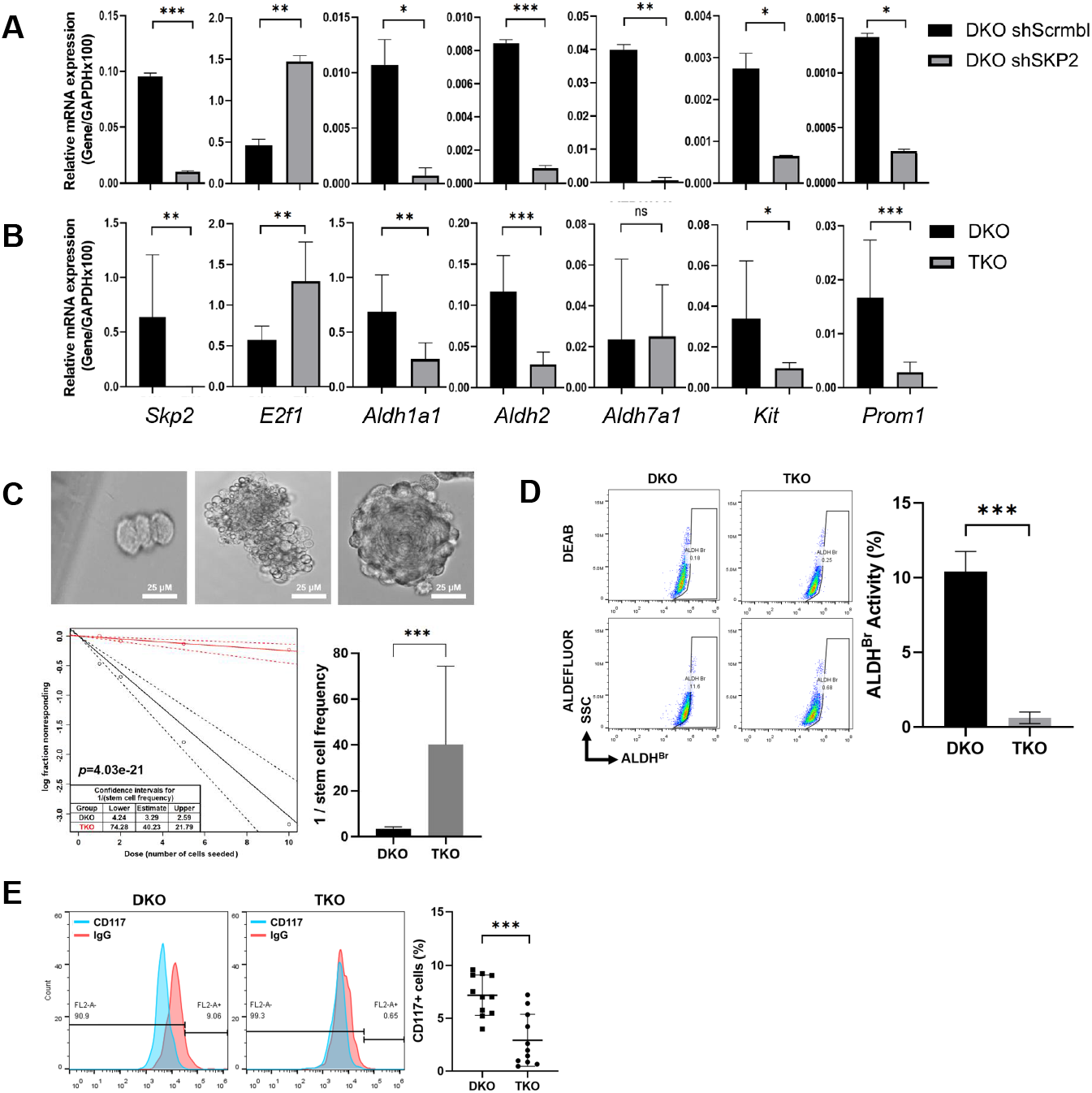
SKP2 deletion inhibits stem-like features of DKO OS. (A) mRNA levels of *SKP2, E2f1, Aldh1a1, Aldh2, Aldh7a1, Kit and Prom1* in DKO cells relative to GAPDH were determined by qRT-PCR followed by SKP2 knockdown (SKO shSKP2). (B) Same genes were also compared among DKO and TKO tumor cells by qRT-PCR to determine mRNA levels relative to GAPDH. (C) Extreme limiting dilution analysis (ELDA) for assessing stem-like cell self-renewal ability. Representative figures were showing cell clusters but not sphere formation (Upper left); a cluster of differentiated and apoptotic cells showing no indication of sphere formation (Upper Middle), and a single sphere formation and scored as “positive” (Upper Right). Scale bars indicate 25 μm. The lower left figure showed sphere-forming capacity output using the previously described algorithm. Graphical representation in the lower right showed the reciprocal of stem cell frequency calculated. (D) ALDH activity staining of tumor cells of indicated genotypes. FACS (representative of three experiments) was used to detect and quantify the subpopulation that expresses a high level of ALDH (ALDH^BR^). (E) Both DKO and TKO tumor cells were stained with CD117, (blue peak) or with the corresponding IgG control (red peak) antibodies. Flow cytometry was performed, and IgG staining was used for gating. Representative results of three experiments were shown on the left, and a quantification comparison of three experiments was on the right. Error bars are SEM. ∗P< 0.05, ∗∗P< 0.01, and ∗∗∗P< 0.001.

### 3.5 Small-molecule inhibitors of SKP2 have anti-tumor activities *in vitro, in vivo*, and in organoids

To examine the efficacy of SCF^SKP2^ inhibitors on OS, we performed MTT assays on DKO, DKOAA and TKO cells using two known SKP2 inhibitors: C1 and Pevonedistat. Mouse osteoblasts (mOB) were extracted from the same mouse strain and served as normal control. Our results indicated that both compounds exhibited strong anti-proliferative effects on DKO cells. In contrast, C1 and Pevonedistat exerted minimal inhibitory effects on TKO and mOB cells (Figure 5A-B), suggesting that these drugs are selective for SKP2-expressing tumor cells. Immunoblotting showed that both C1 and Pevenodistat stabilize p21 and p27 levels in a dose-dependent manner (Figure 5C-D).

**Figure 5.**
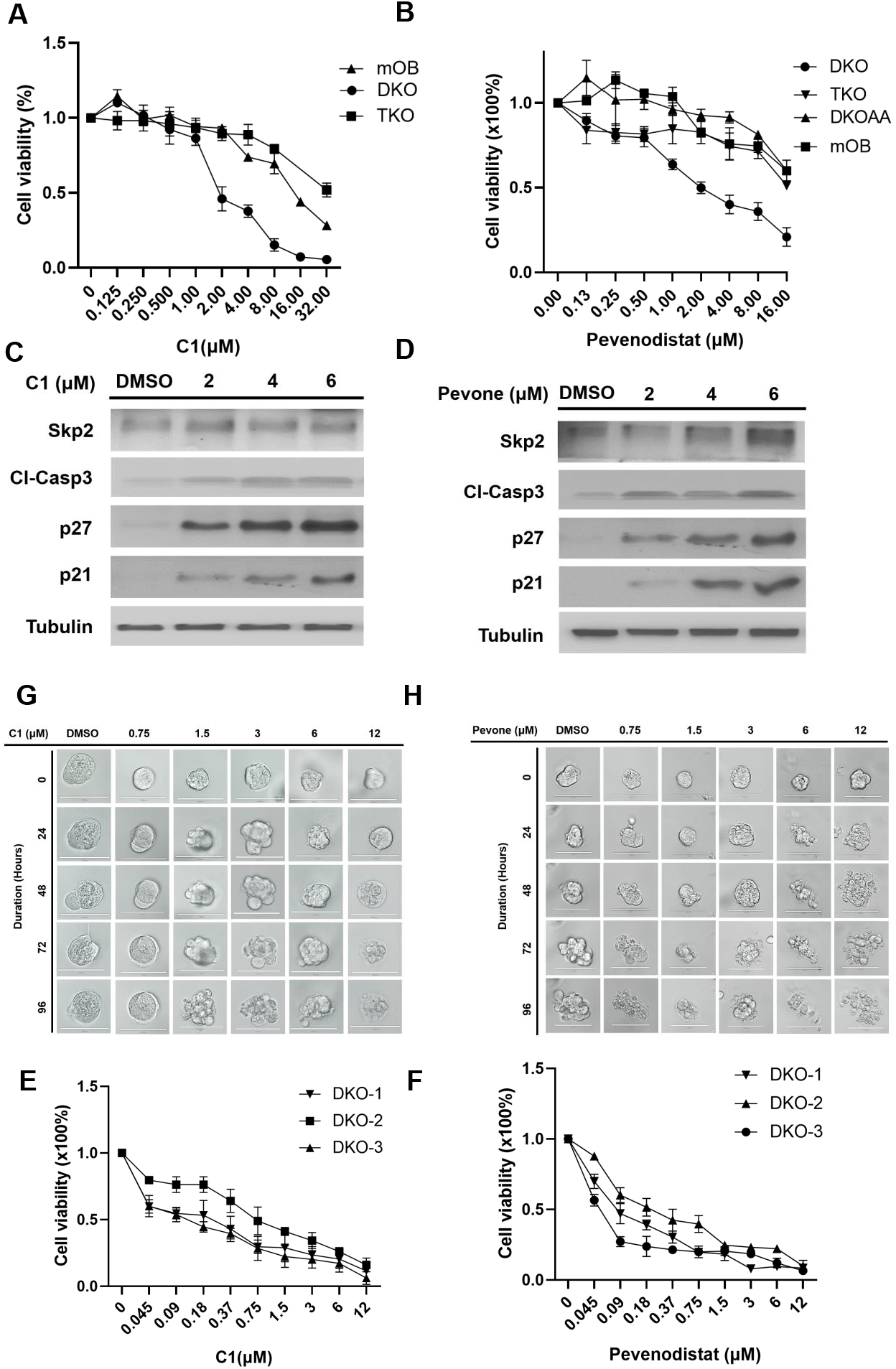
Inhibitors of SCF^SKP2^ suppress DKO tumor cells and tumor-derived organoids. DKO and TKO cell viability following 72 hours of treatment with serial dilutions of SCF^SKP2^ inhibitors, C1 (A) or Penvenodistat (B) at indicated concentrations. Mice osteoblast (mOB) was used as the control. The cell viability was calculated relative to the treatment with DMSO (vehicle control) by the MTT method. Western blot of p21, p27 and cleaved caspase3 in DKO mOS, followed the treatment of C1 (C) or Penvenodistat (D) with indicated concentrations. (E-F) Three independent DKO tumor-derived organoids (DKO-1, DKO-2, and DKO-3) were treated with C1 (E) or Penvenodistat (F) for 96 hours with indicated concentrations. The organoid viability was determined by the CellTiter-Glo method. (G-H) Representative photographs of organoid death and degradation after C1 (G) or Penvenodistat (H) treatment at indicated concentrations and duration. Error bars are SEM.

Next, the effects of SKP2 inhibitors were examined in 3D organoid cultures. We observed growth inhibition of 3 individual DKO organoids in both C1 and Pevonedistat treated groups, as measured by the Glo-Titer method (Figure 5E-H). DKO organoids formed debris piles in response to increased drug concentration and duration of treatment (Figure 5G-H). In a xenograft model, DKO tumor volumes were significantly smaller after daily treatments with C1 (p=0.0084) or Pevenodistat (p=0.0037) (Figure 6A-B). Over one month, body weights did not change significantly at different time points (Figure 6C-D). Consistent with our *in vitro* results, western blots of 4 independent tumor lysates and IHC stainings showed upregulated p27 after drug treatment (Figure 6E-F). The expression of PCNA after drug treatments also mimics the expression seen in TKO tumors (Figure 6G). Together, these results suggest that SKP2 inhibitors exerted a consistent anti-tumor activity in DKO OS.

**Figure 6.**
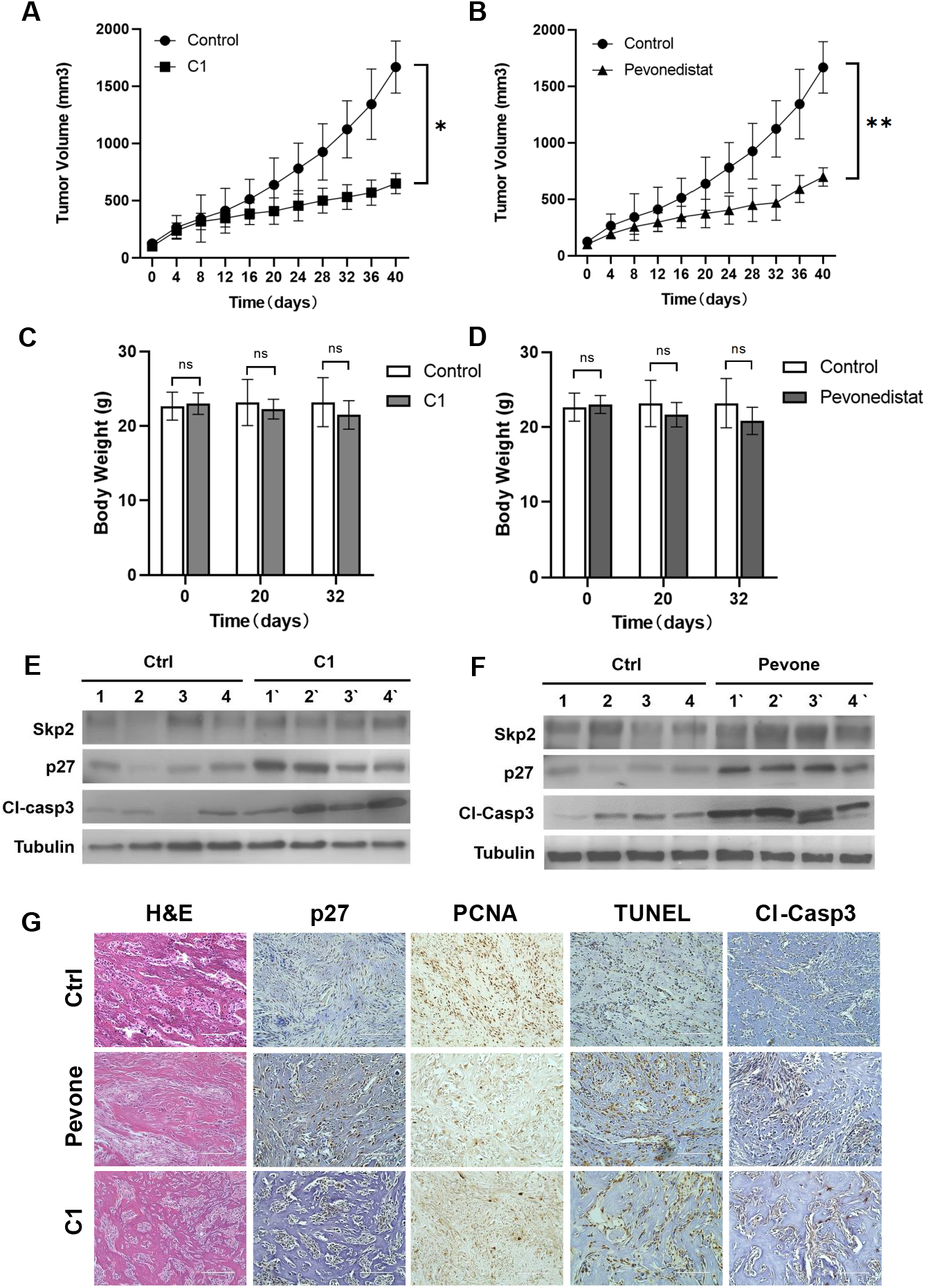
DKO tumors are sensitive to SCF^SKP2^ inhibitors. Tumors were initiated by the subcutaneous inoculation of DKO tumor-derived single-cell suspension. When tumors reached 100 mm^3^ in size, mice were treated by subcutaneous administration of C1 (A; 40 mg/kg/day, *P=0*.*009*) or Penvenodistat (B; 90 mg/kg/day, *P=0*.*022*). The solvent for each drug was used as vehicle control. Tumor size was measured on the indicated days; data are means ± SEM of at least five mice per group. Mouse body weight during daily treatment with C1(C) (*P=0*.*664, 0*.*245 and 0*.*095 at day 0, 20 and 32, respectively*) or Pevonedistat (D) (*P=0*.*693, 0*.*467 and 0*.*220 at day 0, 20 and 32, respectively*), as indicated. (E & F) Western blot and quantitation of SKP2, p27 and cleaved caspase 3 in DKO tumor extracts from control and C1-or Pevonedistat-treated mice. Each lane in (E & F) is an individual mouse tumor. (G) Tumors isolated from drug-treated mice and control were analyzed for the expression of p27, PCNA, cleaved caspase 3 and TUNEL by immunohistochemistry. Statistical significance is indicated by *P < .05,**P < .01, ***P < .001. Column: mean; Error bars are SEM.

### 3.6 The treatment with SKP2 inhibitors promotes apoptosis

To examine the anti-proliferative mechanism induced by SKP2 inhibitors, immunoblotting was performed after the treatment of DKO tumor cells with these drugs. With increasing doses of C1 and Pevenodistat, cleaved caspase-3 was found to increase in a dose-dependent manner, accompanied by an increase in p21 and p27 (Figure 5C, D). To further validate that SKP2 inhibitors induced apoptosis, four xenograft mOS tumors were randomly selected from each group (control, C1, Pevonsdistat) and analyzed. By immunoblotting, cleaved caspase-3 level was consistently elevated in C1 and Pevenodistat treated tumors (Figure 6E-F). Similarly, both TUNEL and cleaved caspase-3 were elevated by IHC in C1 and Pevenodistat treated tumors compared to vehicle control (Figure 6G). This result supports the genetic comparison between DKO and TKO mOS tumors. Conclusively, these suggest that SKP2 inhibitors acted as an effective pharmaceutical method that induces apoptosis to mimic the mechanism in the genetic SKP2 knockout transgenic model and serve as an effective treatment option in mOS.

### 3.7 Small-molecule SKP2 inhibitors act synergistically with conventional chemotherapy for OS

Since we previously showed that C1 worked synergistically with a cytotoxic agent in synovial sarcoma cells[22], we tested whether SKP2 inhibitors have a synergistic effect with doxorubicin, a conventional chemotherapy commonly used in OS. DKO tumor cells were treated with either Pevonedistat or C1 at various doses and combinations based on methods by SynergyFinder 2.0. The cell viability was determined by MTT assays. Both Pevonedistat and C1 showed a strong synergistic effect with doxorubicin, with an average ZIP synergy score of 9.349±1.24 and 6.72±1.28 for Pevenodistat and C1, respectively (Figure 7A-B).

**Figure 7.**
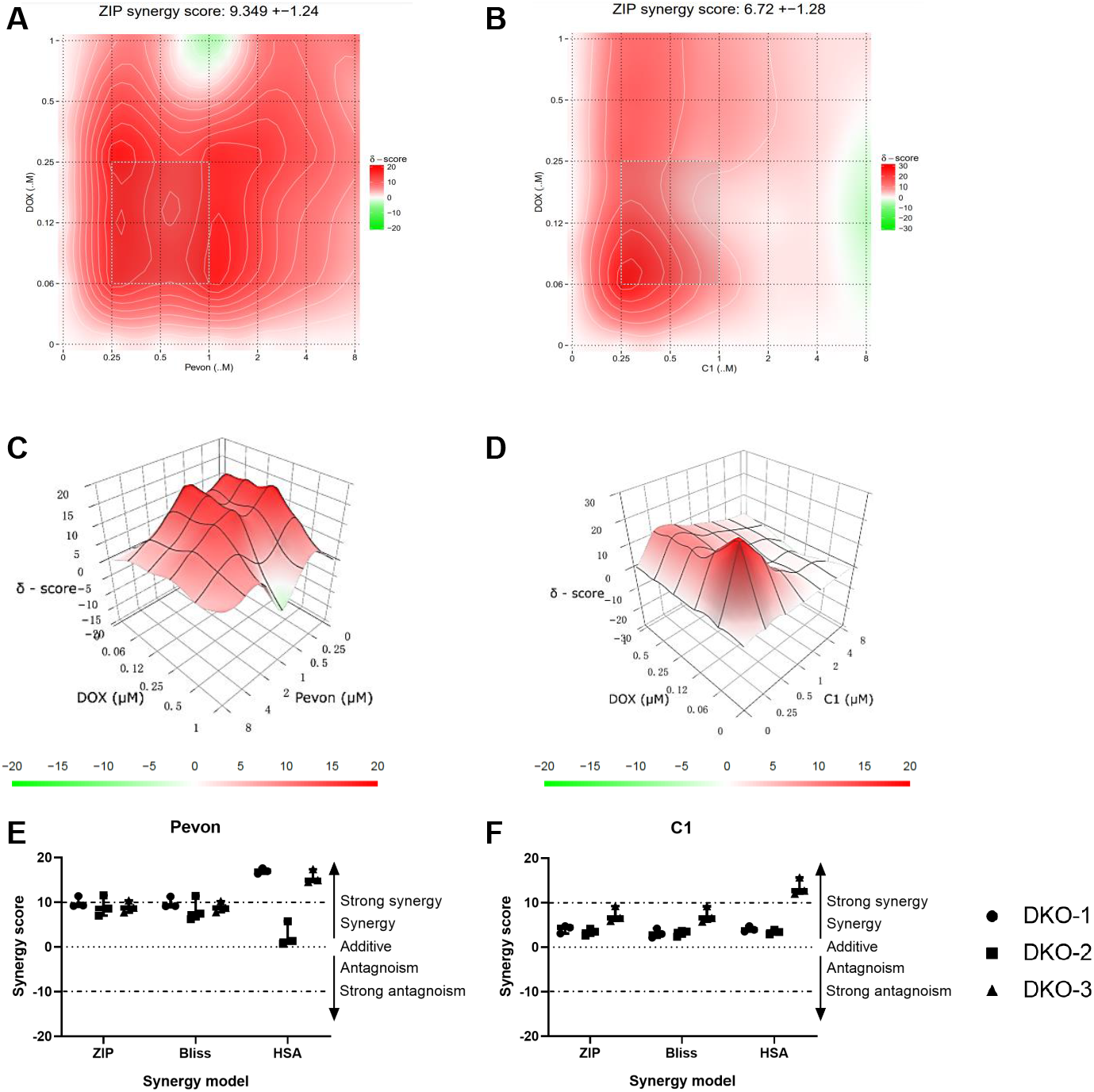
SCF^SKP2^ inhibitors synergize with conventional chemotherapy in DKO OS cells. Analyses of drug combinations between C1 or Pevonedistat with doxorubicin on cell viability were determined by MTT method. The synergistic effects between Pevonedistat and doxorubicin (A), or C1 and doxorubicin (B) were shown in a two-dimensional topographic map of ZIP synergy scores determined by the Synergyfinder. The synergy map highlights synergistic and antagonistic dose regions in red and green colors, respectively. A gray box represents the area with the highest synergy score. (C-D) Synergy landscape determined by ZIP model over C1 or Pevonedistat and doxorubicin concentrations of one representative experiment. Concentrations of C1 or Pevonedistat and doxorubicin as in Spl Fig. S. (E-F) Dot plot of average synergy scores was determined by three different synergy models (ZIP, Bliss and HSA) in three independent DKO primary cultures and repeated three times. Each dot represents one experiment result.

The most synergistic combination was further plotted for each SKP2 inhibitor (Figure 7C-D). To validate the synergistic effect of each drug combination, three different algorithms (ZIP, Bliss, and HAS) were used simultaneously in the analysis of three independent DKO mOS models. Although a non-significant discrepancy can be seen, consistently strong synergistic effects were found with all algorithms (Figure 7E-F).

## 4 Discussion

Although OS is a genetically complex cancer that lacks a defining chromosomal alteration or gene mutation, the pathogenesis of OS is increasingly linked to the inactivation of *TP53*, often potentiated by the loss of function of *RB1* [6, 35] [36]. By re-creating the loss of these tumor suppressors within an osteoblastic lineage, our DKO GEMM recapitulates common genetic and clinical features of the human OS and permits an in-depth inquiry into the role of the ubiquitin degradation pathway in OS tumorigenesis. Previous studies have shown that the F-box protein SKP2 is overexpressed in OS and that *TP53* and *RB1* represent the most common pair of co-inactivated genes among several OS oncogenic drivers [11, 18]. Using data from the PPTC cohort, we found that the expressions of *TP53/RB1* and *SKP2* in OS were inversely correlated in a majority of cases. As a corollary, *SKP2* mRNA and protein levels are also elevated in our DKO model. In addition, deletion of *SKP2* promotes apoptosis, suppresses stem-like properties, and significantly delays the progression of DKO mice to lethality. Taken together, our data support the hypothesis that SKP2 plays an essential role in OS tumorigenesis and suggest that cancers with combined alterations of *Rb1* and *TP53*, such as OS, are particularly vulnerable to SCF^SKP2^ inhibition.

As a tumor suppressor, pRb inhibits SKP2 activities through multiple pathways, involving both transcriptional and post-translational mechanisms [37, 38]. pRb is known to promote SKP2 protein degradation through CDH1 and represses SKP2 mRNA expression via E2F [38, 39]. In addition, pRb can bind directly to the N-terminus of SKP2 and stabilizes p27 by repressing the SKP2 ligase activity [37]. Previous studies have reported that the loss of *SKP2* was sufficient to prevent tumorigenesis in *Rb1*-deficient mice, supporting a dependent role for *SKP2* in *Rb1*-inactivated cancers[30]. In the DKO model, the expression of p27 was found to be reduced in the setting of Trp53 and Rb1 co-deletion. However, levels of p27 were restored after SKP2 was deleted in TKO tumors, which applied to the post-translational role of pRb. Nonetheless, SCFSKP2-mediated ubiquitination and degradation of p27 have been shown as the principal mechanism that impedes the function of p27 as a cell cycle regulator[10]. Consistent with this mechanism, our flow cytometric analyses revealed that TKO tumor cells continued accumulating DNA without dividing, indicating increased cell cycle inhibitors like p27 holding tumor cells from entering phase M in the circumstance without SKP2. The concequence also include the increase of cells in senescence and apoptosis, resulting in a reduced proliferative state.

The transcription factor *E2f1* was consistently elevated when *SKP2* was either deleted in our transgenic model or knocked down by shRNA. This was followed by a corresponding upregulation of E2f1 target genes, including *Bbc3, Bid, Bcl2l11* and *casp3*. Therefore, inhibiting SKP2 likely promotes a pro-apoptotic microenvironment, which contributes to the delay in tumorigenesis and survival advantage of TKO animals. Interestingly, our analysis of the NCI TARGET database also revealed that apoptotic-related genes are associated with a better prognosis in OS patients. Previous studies have shown that when *SKP2* is co-deleted with *Rb1*, high levels of p27 sequester Cyclin A and prevent its binding to the E2f1 promoter. This activates E2f1 target genes and converts E2F1 from an oncogenic to a pro-apoptotic factor[30, 40]. Finally, our results are consistent with previous studies showing a synthetic lethal relationship between Rb1 and SKP2 deletions[41], providing a strong rationale for the use of SKP2 inhibitors in treating Rb1-inactivated OS.

In our study, the survival advantage of TKO mice suggests that targeting the SCF^SKP2^ complex may have clinical relevance. Since Pevonedistat and C1 showed more efficacy in DKO cells than in TKO cells or normal osteoblasts, these compounds likely have a degree of selectivity for SKP2-expressing tumors. Our *in vivo* testings also confirmed that these compounds exerted a consistent anti-tumor effect in DKO OS xenografts with a minimal toxicity. At present, Pevonedistat is the only neddylation inhibitor of SCF^SKP2^ complex in human clinical trials [42, 43]. In the future, as more SCF inhibitors become available, we will likely need high-throughput preclinical studies using 3D organoid technologies or patient-derived xenografts to assess the efficacies of these new compounds to reduce the OS disease burden.

Genetic and pharmacological inactivation of SKP2 has been shown to reduce the CSC population in prostate cancer [23]. Using the DKOAA transgenic model, we recently showed that the binding between SKP2 and p27 likely enhances OS cancer stemness[18]. In the current study, TKO cells formed fewer tumorspheres and exhibited a lower ALDH activity and self-renewal capacity compared to DKO cells, suggesting that SKP2 plays an essential role in maintaining tumor-initiating properties in OS. Therefore, inhibitors of the SKP2-p27 axis may have a therapeutic utility by targeting the CSC subpopulation. Previous studies have shown that CD117 is preferentially expressed in tumorspheres and chemo-resistant OS cells [44]. Here, we found that an SKP2 deletion significantly reduced the CD117 subpopulation in OS tumors, suggesting that SKP2 promotes a chemo-resistant phenotype. Taken together, our results raise the possibility that adding SKP2 inhibitors into the current chemo regimen for OS can improve outcomes by targeting the cancer stem cell population and reducing chemoresistance in OS.

In our study, the deletion of *SKP2* markedly prolonged the survival of DKO mice but did not block tumorigenesis entirely. Given a high degree of genomic instability in OS, it is conceivable that secondary genetic alterations or alternative pathways may provide an escape mechanism in response to an SKP2 deletion. On the other hand, the difference in survival between TKO and DKOAA mice suggests that the oncogenic functions of SKP2 may extend beyond its canonical role in degrading p27. For example, the role of SKP2 in ubiquitinating another substrate such as p21 may also play a role in OS progression. However, given that p53 is inactivated in our models and p21 is a known downstream effector of p53, one may argue that the contribution of p21 may be less consequential in DKO tumors. Cks1 is an adapter protein that facilitates the interaction between SKP2 and p21/p27 docking. In the future, generating a transgenic model to examine the role of Cks1 may be desirable to understand the pathobiology of OS[45].

The low success rate of bench-to-bedside drug development suggests a lack of suitable models that can replicate the tumor microenvironment with high fidelity. More recently, organoid culture technologies have been developed as an intermediary between *in vitro* cancer cell lines and patient-derived xenografts[46]. Even after a long-term expansion, cancer cells grown in 3D organoid cultures retain the architecture, morphology, and cell-matrix interactions that closely resemble those of the original tumors, as demonstrated in prostate, pancreatic, and lung cancers [10, 16, 47, 48]. However, most organoid models have been generated from epithelial cancers yet are rarely reported in mesenchymal cancers[49]. Here, we established a reproducible culture system to generate OS organoids, which can be employed for drug testing within a short time frame. Our histological analysis showed a similarity between the original tumor tissue and the organoid culture model. Our organoid cultures represent the first step toward replicating the genetic diversity of OS and serve as a proof of principle to translate our mouse model findings into patient-specific treatment. In the future, it is conceivable that a personalized, high-throughput drug screening system can be established from patient-derived OS organoids, which offer a unique opportunity to identify effective SKP2-related therapies for individual patients.

## Conclusion

In this study, we found that the co-inactivation of *RB1* and *TP53* in OS often correlates with an increase in *SKP2* expression. In addition, our data reveal the following key findings: (1) targeting SKP2 promotes apoptosis, cell cycle arrest, and inhibits cancer stemness, contributing to a survival benefit in DKO OS; (2) Small-molecule SKP2 inhibitors are effective in a preclinical setting against Rb/p53-deficient OS and exhibit synergy with conventional chemotherapy. Therefore, combination trials of SKP2 inhibitors and conventional chemotherapy may be desirable to overcome the stemness plasticity of OS tumors and therefore minimize the risk of relapsed disease. In the future, efforts to decipher intrinsic functions of other components of the SCF complex may lead to more specific approaches to improve OS survival.

## Supporting information

supplemental meterial

## Competing interests

All authors declare no conflict of interest.

## Acknowledgments

J.W., A.F., S.S., R.Z., and H.Z. participated in the study design, performed the study, and prepared the manuscript. J.W., A.F., S.S., R.Z., V.V., W.A., O.A., H.B., A.S., J.T., and S.Y. participated in the study design and performed experiments. A.F., D.Z., Y.L., and J.W. performed statistical and bioinformatic analysis. R.G., X.Z., D.Z., E.L.S., H.Z., R.Y., D.S.G., D.Z., and B.H.H. conceived the study, participated in the design and coordination, and manuscript editing. Fundings were provided by grants from Sarcoma Strong, Montefiore Medical Center (to B.H.H.), and the National Institutes of Health (R01CA255643 to B.H.H; R01CA193967 to X.Z.; R01CA230032 and R01201458 to E.S. and H.Z.) and the National Natural Science Foundation of China (82103223 to J.W.).

## Notes

### Competing Interest Statement

The authors have declared no competing interest.

