## supplemental meterial for "Targeted inhibition of SCF^SKP2^ confers anti-tumor activities resulting in a survival benefit in osteosarcoma"

### A targeted inhibition of $SCF^{SKP2}$ has anti-tumor activities that result in a survival benefit in osteosarcoma

#### Supplementary Tables

**Table S1** Genotyping primers used for identification of Skp2 knockout transgenic mice
**Table S2** qPCR primers used in this study
**Table S3** Table showing results of survival analysis of the Reactome Apoptosis gene signature in the NCI TARGET OS cohort

#### Supplementary Figures

##### Figure S1

Genotyping results using agarose gel electrophoresis. (A) Genotyping results of *Trp53* wild type(*p53*$^{(+/+)}$)(288bp), p53-lox heterozygotes (*p53*$^{(lox/+)}$) (288bp and 370bp) and p53-lox homozygotes (*p53*$^{(lox/lox)}$) (370bp). (B) Genotyping results of Osterix1-Cre (*Osx-Cre(+)*) (198bp) and coresponding internal positive control（*Osx-Cre(+)* Ctrl) (253bp). (C) Genotyping results of *Rb1* wild type (*Rb1*$^{(+/+)}$)(250bp), Rb1-lox heterozygotes (*Rb1*$^{(lox/+)}$) (250bp and 310bp) and Rb1-lox homozygotes (*Rb1*$^{(lox/lox)}$) (310bp). (D) Genotyping results of recombined *Trp53* and *Rb1* after deletion mediated by Osterix-Cre. ( ~ 500 bp). (E) Genotyping results of wild type *p27* (250bp), *p27 T187A* heterozygotes (*p27*$^{(T187A/+)}$) (250bp and 284bp) and T187A homozygotes (*p27*$^{(T187A/T187A)}$) (284bp). (F) Genotyping results of wild type *Skp2* (*Skp2*$^{(wt/wt)}$) (500bp only) and *Skp2 knockout* homozygotes (*Skp2*$^{(-/-)}$) (430bp only).

##### Figure S2

Sashimi plots showing splicing RNA-seq reads mapped to the *Trp53* (A), *RB1* (B) and Skp2 loci(C). The numbers above the arch lines indicate the numbers of reads for individual splicing patterns. (C) Summary of RNA-seq reads mapped to the T187A mutation sites in the DKO (top) and DKOAA (bottom), confirming mutation in DKOAA samples.

##### Figure S3

Kaplan-Meier survival analysis comparing the DKO, DKOAA and TKO cohorts of mice undergoing tumorigenesis exclude maxillofacial tumor. P-value is by a log-rank test and indicated in the figure.

##### Figure S4

Graph showing the tumor anatomical distribution of each genotype.

##### Figure S5

Mouse body weight during daily treatment of 40 days with either C1 or Pevonedistat as indicated. Statistical significance is indicated by $^*P < .05$,$^{**}P < .01$, $^{***}P < .001$. Column: mean; Error bars are SEM.

**Table S1 Genotyping primers used for identification of Skp2 knockout transgenic mice**

| | | |
|---|---|---|
| Wildtype allele | KN3 | 5′- AGAGTGGAAGAACCCAGGCAGGAC-3’ |
| | KN4 | 5′- CCCGTGGAGGGAAAAAGAGGGACG-3’ |
| Knockout allele | KN13 | 5′- GCATCGCCTTCTATCGCCTTCTTG-3’ |
| | KN38 | 5′- TTCCCACCCCCACATCCAGTCATT-3’ |

**Table S2 qPCR primers used in this study**

| | | |
|---|---|---|
| CDKN1B | sense | 5′-GCGGTGCCTTTAATTGGGTCT |
| | antisense | 5′-GGCTTCTTGGGCGTCTGC T |
| p73 | sense | 5′-AACGCCGAGCATCAATCC |
| | antisense | 5′-AGCCCAGACTCTGAGCACTT |
| Skp2 | sense | 5′-AGCAGCCGCTGGGTGAAAGC |
| | antisense | 5′-ATCACTGAGTTCGACAGGTCCAT |
| E2f1 | sense | 5′-TCACTAAATCTGACCACCAAACG |
| | antisense | 5′-TTGGACTTCTTGGCAATGAGC |
| Bbc3(Puma) | sense | 5′-GGTCCAGACTGTGAATCCTGTG |
| | antisense | 5′-TCCTCCCTCTTCTGAGACTTCC |
| Bid | sense | 5′-ACGGAATGCAAAGAACAACTC |
| | antisense | 5′-CAACGCTTGAGGATACAGTGAG |
| Bcl2l11(Bim) | sense | 5′-ATCTTGTTGGGCTTACTTGTG |
| | antisense | 5′-GTCCTGCCTGGTCTTGAAAT |
| Casp3 | sense | 5′-CGCGCACAAGCTAGAATTTATG |
| | antisense | 5′-GGACACAATACACGGGATCTG |
| Prominin1 | sense | 5′- TCTGCTGACATTTGCCTCTAC |
| | antisense | 5′- GCTGGTGGATGGCTCTTATATT |
| ALDH1A1 | sense | 5′- GAGAGTGGGAAGAAAGAAGGAG |
| | antisense | 5′- CTCATCAGTCACGTTGGAGAA |
| ALDH2 | sense | 5′- CATCTTGGTACCTGGGATCTTG |
| | antisense | 5′- TGTAGCTGCAGCCAAGAATAG |
| ALDH7A1 | sense | 5′- CTCAGTACCACCACAACAAAGA |
| | antisense | 5′- CAAGAACGAAGGTCTGTCTACC |
| CD117(Kit) | sense | 5′- AGGAGAACTGAGGCTGTTTG |
| | antisense | 5′- TAACTTGTGCTCCCTGCTATG |
| GAPDH | sense | 5′- GGTTGTCTCCTGCGACTTCA |
| | antisense | 5′-GGTGGTCCAGGGTTTCTTAC |

**Table S3.**

| | Hazard Ratio | Lower 95% CI | Upper 95% CI | Wald P-value |
|---|---|---|---|---|
| Dichotomized, Hi vs Low | 0.301 | 0.125 | 0.723 | 0.007 |
| Univariable Continuous | 0.964 | 0.932 | 0.996 | 0.028 |
| Multivariable Continuous | 0.968 | 0.939 | 0.999 | 0.04 |

**Figure S1**

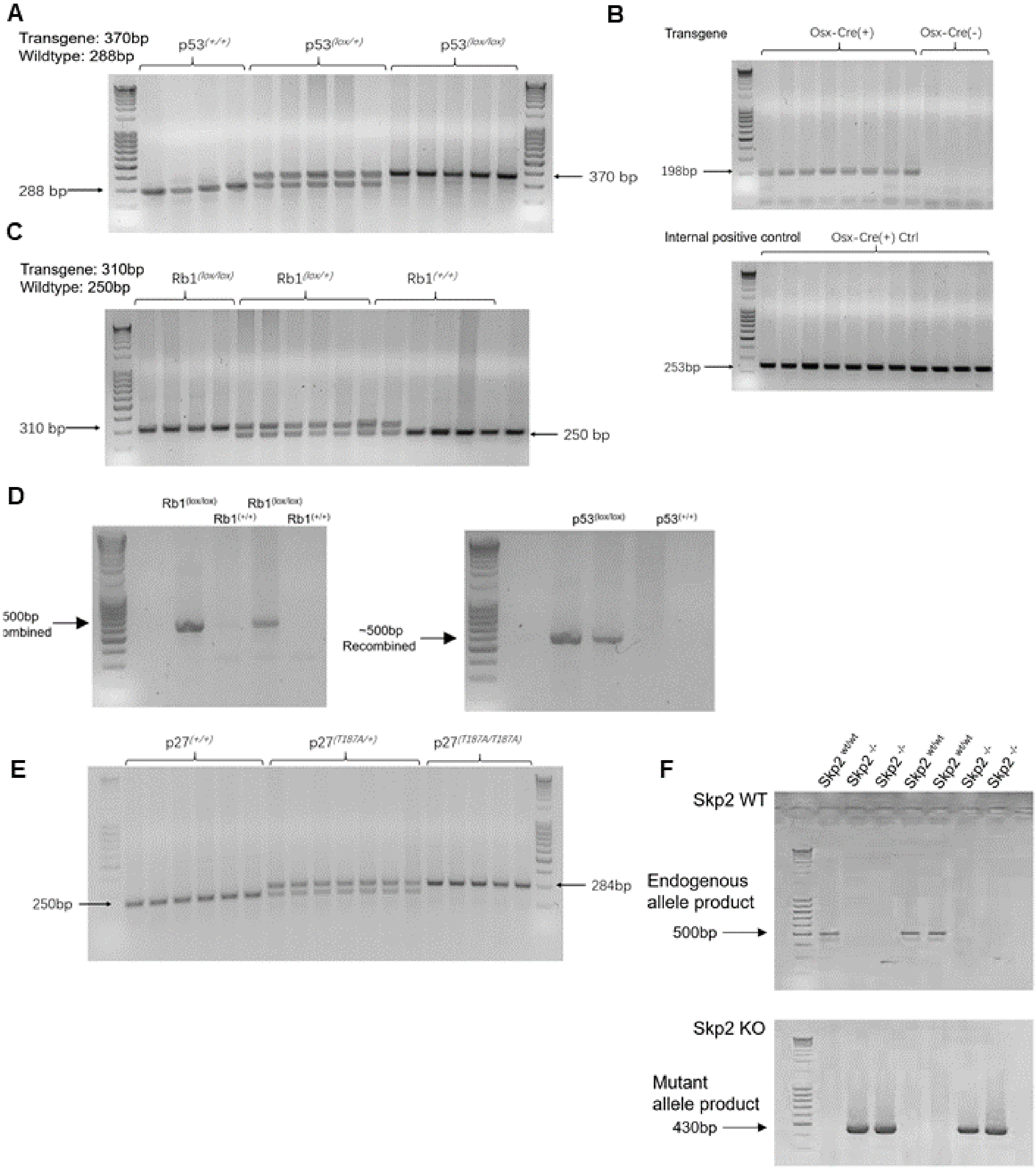
A
Transgene: 370bp
Wildtype: 288bp
p53(+/+)
p53(lox/+)
p53(lox/lox)
288 bp
370 bp
B
Transgene
Osx-Cre(+)
Osx-Cre(-)
198bp
Internal positive control
Osx-Cre(+) Ctrl
253bp
C
Transgene: 310bp
Wildtype: 250bp
Rb1(lox/lox)
Rb1(lox/+)
Rb1(+/+)
310 bp
250 bp
D
Rb1(lox/lox)
Rb1(+/+)
Rb1(lox/lox)
Rb1(+/+)
500bp
ombined
~500bp
Recombined
p53(lox/lox)
p53(+/+)
E
p27(+/+)
p27(T187A/+)
p27(T187A/T187A)
250bp
284bp
F
Skp2 wt/wt
Skp2 -/-
Skp2 -/-
Skp2 wt/wt
Skp2 wt/wt
Skp2 -/-
Skp2 -/-
Skp2 WT
Endogenous
allele product
500bp
Skp2 KO
Mutant
allele product
430bp

#### Figure S2

A) Splicing reads at exons 2-10 excision of Trp53, DKO (up) and TKO (down)

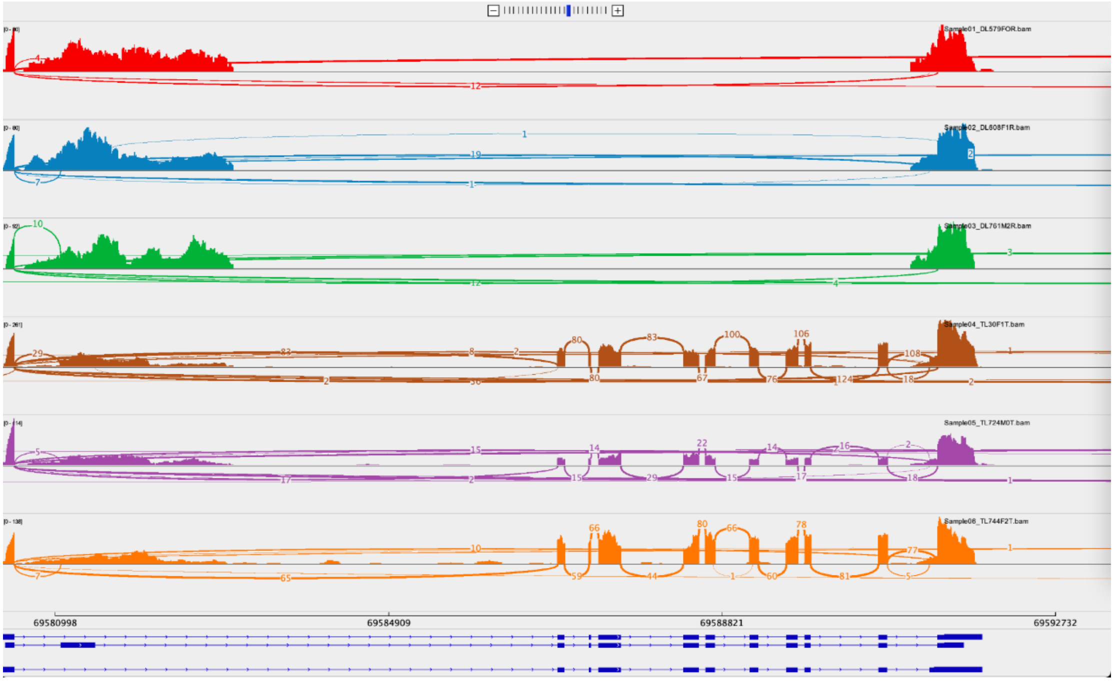


B) Splicing reads at exons 3 excision of RB1, DKO (up) and TKO (down).

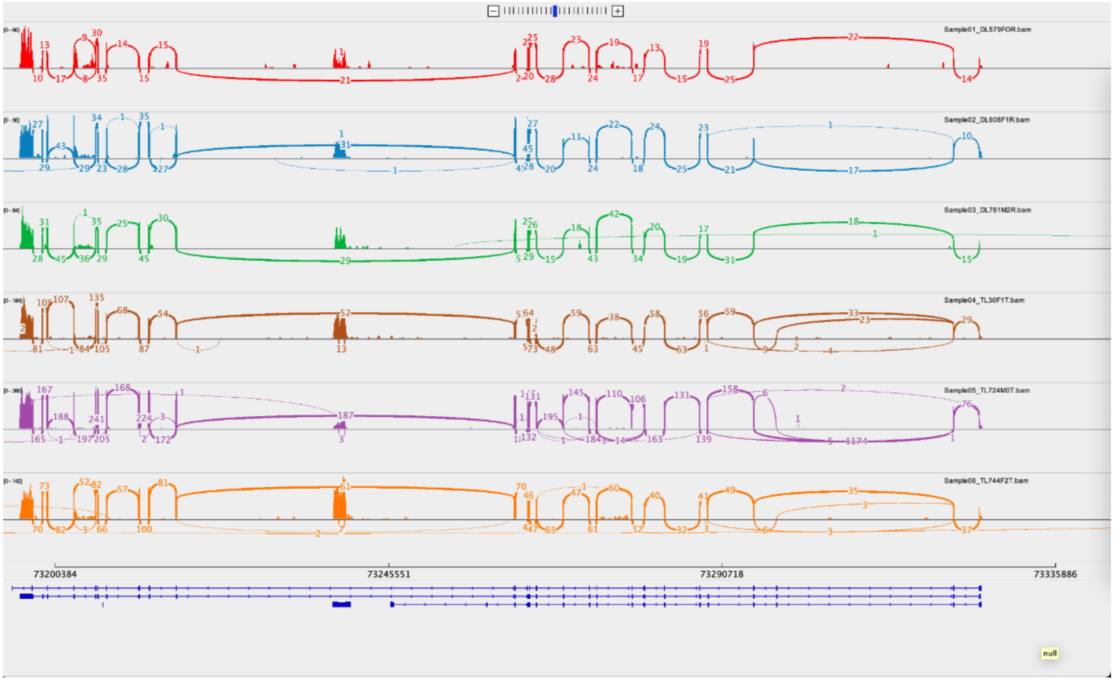

C) Splicing reads showing deletion of Skp2, DKO (up), and TKO (down).

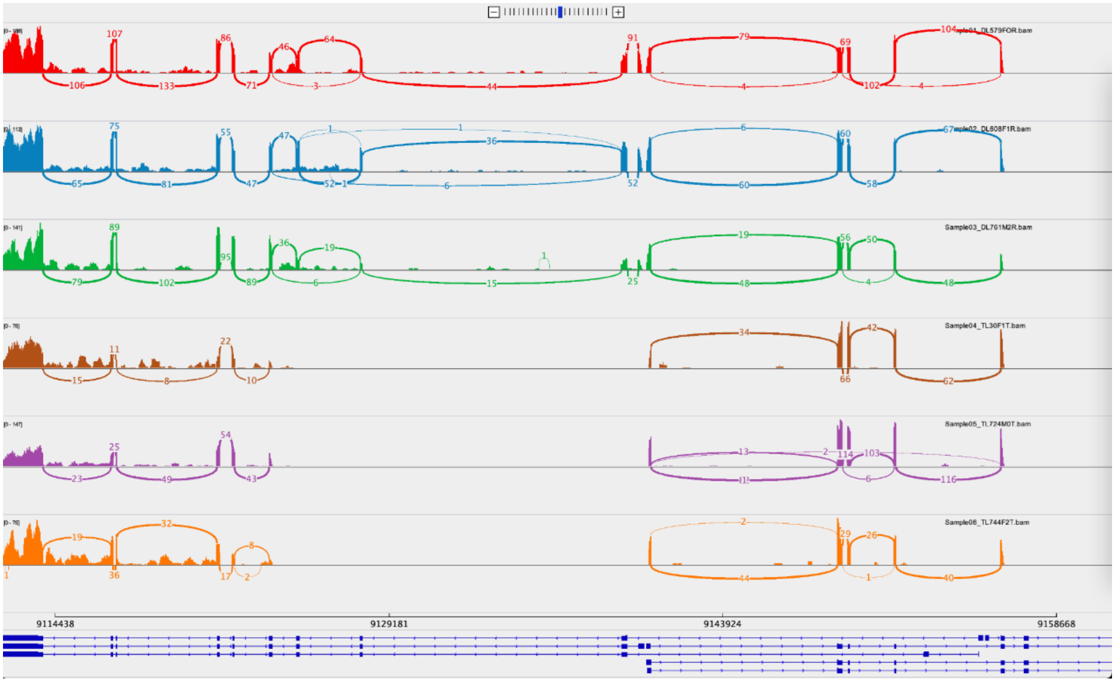


#### Figure S3

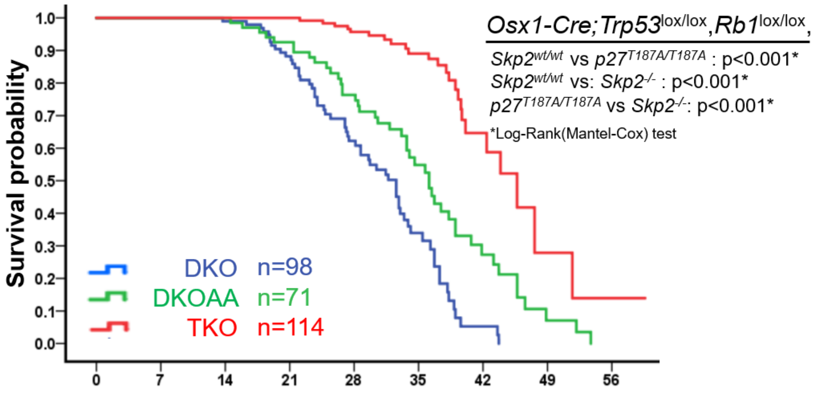

#### Figure S4

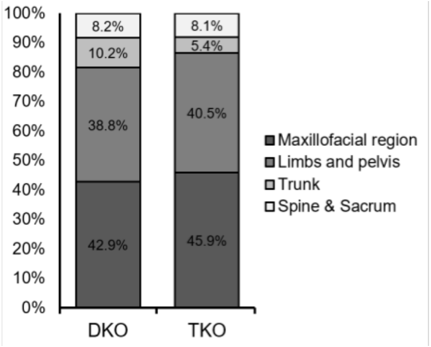


#### Figure S5

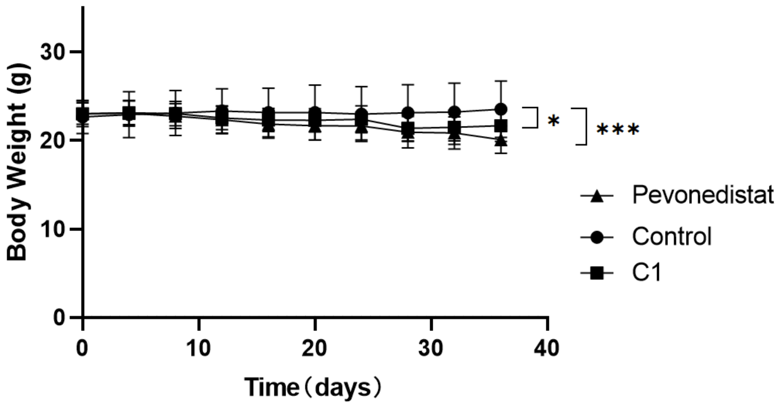
